# Life trajectories towards suicide: developmental role and specificities of adolescence

**DOI:** 10.1101/2020.01.18.911230

**Authors:** Charles-Edouard Notredame, Nadia Chawky, Guy Beauchamp, Guillaume Vaiva, Monique Séguin

## Abstract

**Objective:** Considering that adolescence is a crucial period of allostatic maturation, we aimed at testing whether it represents a breakpoint in adversity trajectories to suicide and, relatedly, whether life trajectories of adolescents who died by suicide differ from those of adult counterparts.

**Method:** In 303 individuals who died by suicide, a panel of experts derived longitudinal burden of adversity ratings from extensive clinical interviews conducted with informants. Piecewise Joint Latent Class Models allowed to identify patterns of adversity trajectories and test the introduction of breakpoints in life-paths. The classes inferred from the optimal model were compared in terms of socio-demographics, psychopathology and nature of experienced events.

**Results:** The most accurate model derived 2 trajectory patterns with a breakpoint in early adolescence. In the first class (n = 43 individuals), the burden of adversity increased steadily from birth to death, which occurred on average at 23 (SE = 1.29). In the second class (n = 259), where individuals died on average at 43 years (SE = 0.96), the burden of adversity followed a similar trajectory during infancy but stabilized between ages 10 and 14 years and rebounded at about 25. The classes differed in terms of childhood family stability, occurrence of dependent events, distal exposure to suicide and intra-family sexual victimization.

**Conclusion:** Although subject to inherent limitations of the retrospective designs, our results encourage clinicians and researchers to pay greater attention to the diathetic maturation processes occurring during adolescence when considering suicidal behaviors in young people.

## 1. Introduction

Among 15- to 29-year-olds, suicide represents the second most common cause of death and accounts for 8.5% of young people dying^1^. Interestingly, suicidal behaviors generally appear, but also peak, immediately after puberty^49^, suggesting that adolescence may be a critical period—both a tipping and a frailty point—in the suicidal process.

Developmental approaches, which focus on how risk factors dynamically integrate at an individual level^4,5^, offer a convenient framework to study how the socio-biological transformations of adolescence and the suicidal process may heuristically shed light on each other. Turecki and Brent proposed to categorize most influential risk factors based on their putative role in the developmental sequence towards suicidal behaviors^6^. The authors described distal risk factors as the early biological and environmental determinants that durably shape a person’s vulnerability to suicide, developmental risk factors as the phenotypic expressions of this vulnerability and proximal risk factors as the clinical conditions or triggering negative life events that contribute to precipitating suicidal behaviors. The model builds upon a stress-diathesis conception of suicide, according to which individuals are self-regulated organisms that adapt to a changing environment in order to maintain their homeostasis^7,8^. When detected as a threat, Adverse Life Experiences (ALE) elicit neuro-biological reactions to maintain the equilibrium^8^. At the behavioral level, allostatic load^9^ (i.e. stress-induced deviation from homeostasis) triggers goal-oriented coping reactions to solve the adverse experience and/or minimize its subjective painful consequences. Some individuals may exhibit an increased probability of dysfunctional coping responses to stress – a condition known as diathesis. Suicide is then understood as the most extreme form of abnormal coping strategy due to stress overload exhausting or overwhelming regulatory mechanisms^9,10^.

Developmental models of suicidal behaviors still crucially lack proof of concept, as traditional epidemiological methods appear ill-suited to the multi-deterministic and interactionist postulates of developmental psychopathology^4^. As a more dynamic alternative, developmental methods are increasingly used in research on suicidal behaviors to retrace the longitudinal evolution of ideations^11–13^, covariates^14^ or predictors^15–17^. For more than 20 years, our team has sought to take the understanding of the causal process towards suicide a step forward by computing trajectories of Burden of Adversity (BA)^5,18,19^. Derived from extensive narrative material, the notion of BA integrates not only the ALE that a person has encountered in his/her life, but also the severity, interactions, repetitions and context of these ALE. Conceptually, the BA is deemed to reflect the allostatic load that weighs on a given individual, but also, and inseparably, the level of the stress-diathesis interaction that characterizes the way this individual responds to adversity.

Incidentally, the stress-diathesis model of suicide gives a glimpse of the pivotal role that adolescence may play in trajectories towards suicide. As of adolescence, the model predicts that the functional status of risk factors changes, with adversity undergoing a shorter-term precipitating influence and diathesis stabilizing and being appraisable through developmental risk factors. Implicitly, such a functional switch would reflect a transition in the nature of the stress-diathesis interplay, possibly due to the maturation of the allostatic systems. Along the same line, our group evidenced that the life course of individuals who died by suicide before the age of 30 differed significantly from the trajectory of individuals still alive at that same age^19^. Although offering first clues of the specific vulnerabilities of adolescents and young adults in regard to suicide, these results leave obscure zones to be explored. More specifically, the transitional value of adolescence in trajectories towards suicide remains purely hypothetical and lacks supportive evidence.

The goal of this paper is to explore whether the life trajectories of adolescents who died by suicide differ significantly from those of adults who died by suicide and whether the developmental process involved in adolescence plays a role in these trajectories. To address this dual objective, we advanced the hypothesis that both the developmental role and the specific vulnerability of adolescence as regards suicide paths are related to the same underlying developmental process.

## Methods

### 1.1. Sample and recruitment procedure

Participants were recruited in the provinces of Québec and New Brunswick, Canada. The sample came from 4 successive recruitment campaigns conducted between 2003 and 2015. We included all the cases of suicide registered by the provincial Chief Coroner’s office during the corresponding periods. On average, 75% of identified cases were included. Reasons for non-inclusion were over-riding legal contingencies, absence of informant, or dissent within the family. Upon agreement of the relatives, the research team inquired about the person who had best known the deceased to propose him/her appointments as the referent informant.

The protocol received approval from the ethic review boards of the Douglas Mental Health University Institute, the Centre Hospitalier Universitaire Sainte-Justine and the Université du Québec en Outaouais. All informants signed a consent form.

### 1.2. Data collection

#### 1.2.1. General procedure

Skilled investigators conducted 2 to 3 in-depth interviews with each informant, focusing on their deceased relative. The interviews occurred between 6 and 18 months after the death, lasted 2 to 3 hours on average and comprised three sections: exploration of sociodemographic characteristics and medical history, psychopathological investigation and inventory of ALE. The data collected during the interviews were crossed-checked and completed by accessing the deceased’s medical files and social services charts when available.

#### 1.2.2. Psychopathology

Each case was submitted to a post-mortem diagnostic assessment according to the psychological autopsy standards. The investigators administered the Structured Clinical Interview for DSM-IV for Axis I and Axis II disorders (SCID-I and SCID-II) to the informant, who was invited to respond in reference to his/her relative. A panel of experts reviewed the resulting material to determine, by consensus, the presence of a diagnosis 6 months prior to death and lifelong. “By-proxy” diagnostic procedures have proven good to excellent reliability and strong concordance with directly administered diagnostic interviews^15,16,22,23^.

#### 1.2.3. Inventory of adverse life experiences

To collect all the possible ALE that the subjects had encountered, we adopted narrative methods borrowed from life history calendar research^20^. This investigation technique has been described in detail elsewhere^18^. In brief, it consisted of a semi-structured conversational exploration designed to maximize the chances of identifying significant life events. The narrative screening was conducted along a double axis of progression: chronological (i.e. from birth to death) and dimensional, as we systematically explored 9 pre-defined spheres of life. The retrospective recall was guided by visual timelines where informants pinpointed memory anchors such as significant biographical elements. The length, frequency, severity and context of each reported event were systematically collected. All the interviews were recorded.

### 1.3. Data transformation

Qualitative data collected from the narrative interviews were quantitatively transformed according to a human-rating procedure. After each interview, the investigators drafted a synthetic clinical vignette out of the subjects’ life calendar, which was then submitted to a panel of independent expert raters. In a clinical decision process, the raters were asked to integrate, for 5-year periods, all the ALE that had occurred, their circumstantial and developmental context and the history of the individuals to estimate the average “contextual threat” that had been weighing on them. This estimation took the form of a BA value ranging from 0 (low) to 5 (severe) based on standard definitions for each level of BA. The experts rated each trajectory independently before a consensus discussion.

### 1.4. Data analysis

After computing individual BA evolution, we used Joint Latent Class Modeling (JLCM) to derive typical adversity trajectories while accounting for the time-dependent risk of dying^21,22^. Each class was specified by its own latent growth parameters, estimated from the BA variance-covariance matrix, and the hazard function parameters, derived from the corresponding survival data set. The class membership probability was estimated over both the processes.

To test whether adolescence represented a breaking point in the developmental trajectories to suicide, we then computed piecewise JLCM where the value of the latent slope (± quadratic parameter) in each class was authorized to change as of a pre-determined timepoint^23^.

The resulting structure we used for our models is represented in Figure 1. We implemented all possible variations of this structure according to the following factorial development: [2, 3 or 4 classes] x [linear or linear + quadratic curve] x [no break, break at 15, 20, or 25]. A quadratic term was added only on curve segments for which the number of points was sufficient for the model to be specified. Class-specific growth and hazard parameters were adjusted on the subjects’ gender and recruitment campaign as time-invariant covariates.

**Figure 1.**
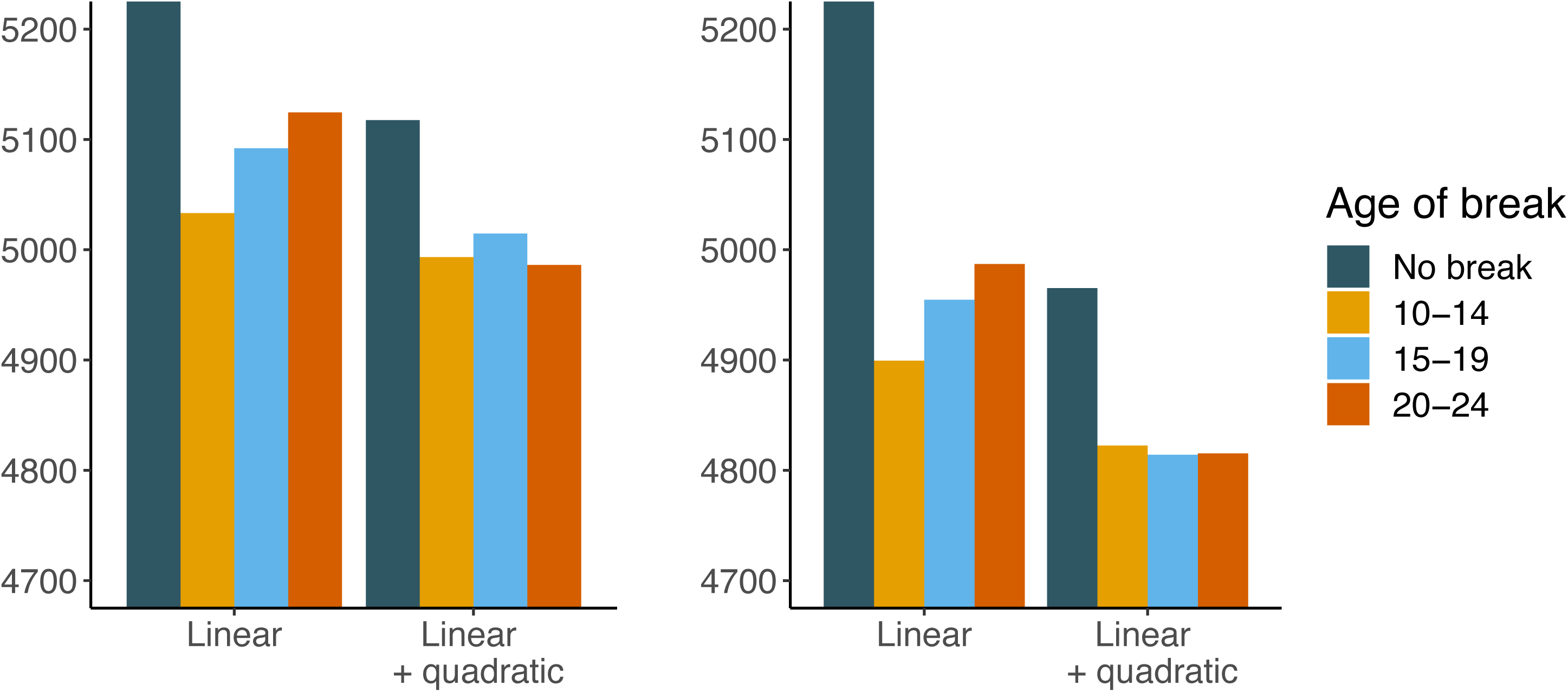
Diagram of the Joint Latent Class Model. Circles represent latent variables or processes: intercept (i), slope (q) and quadratic term (q) of the growth process, hazard function (f) and clustering variable (c). Squares represent observed variables: burden of adversity, survival data and covariates (gender and recruitment campaign). The specification of different successive slopes (s1 and s2) and quadratic terms (q1 and q2) allows piecewise Modeling. The 2 pieces of the trajectories are differentiated by their blue and yellow colors.

Because the proportion of people alive progressively decreased with age, resulting in poorer information, we decided to fit our model on the 7 first time-points of available BA values, i.e. until age 34 years. Remaining non-ignorable missing data due to people dying were taken into account by the JLCM^24^.

Once the best performing model was selected based on fit index, parsimony and clinical relevance criteria, we conducted pairwise intra- and inter-class comparisons of the growth parameter segments using Wald tests. We also compared the 2 classes in terms of sociodemographic characteristics, psychopathology and occurrence of ALE during distal (0-9 years old), proximal (year prior to death) and trajectory break-point periods. We used 2-sided Wilcoxon or Student tests to compare continuous distributions and Chi-square of Fisher tests to compare proportions. The alpha risk was fixed at 0.05.

Statistics were conducted with Mplus Version 7.4^25^ and R version 3.6.1^26^

## 2. Results

Our sample consisted of 303 individuals. Seventy percent were men. Mean age at death was 40.5 (SD. 16.3).

### 2.1. Model predictions

The model we retained as best accounting for our data was a 2-class, quadratic piecewise LCGM with a break at age 10-14. The detailed selection process is described in the Supplemental Material.

The predicted BA trajectories corresponding to each class are represented in Figure 2A. Parameter estimates are presented in Table 1. In the first class, which included 39 individuals (13% of the sample) who died at mean age 23.2 (SD. 8.0), the BA started from a non-null value of 1.25 (SE. 0.07) at birth and steadily increased at a rate of 0.51 (SE. 0.09) to 0.72 units (SE. 0.21) per 5 years until death. The growth was rather smooth and linear, as the quadratic term was non-significant and the Wald test did not reveal any significant difference between the slope parameters of the curve segments. The remaining 264 (87%) individuals of the sample followed a BA trajectory that also increased from birth to young adolescence at a rate of 0.31 (SE. 0.03) units per 5 years. However, the growth in BA significantly dampened at age 10-14 to progressively increase again, as suggested by a non-significant slope of 0.08 (SE. 0.06, comparison with the slope of the first segment: W = 32.69, p < 0.001), but a significant quadratic term of 0.07 (SE. 0.07). In this class, death occurred at mean age 43.1 (SD. 15.6). Interestingly, the intercept (BA at birth) did not differ significantly between the 2 classes (p = 0.85), while the slope of the first segment of the curves did mildly (p = 0.02).

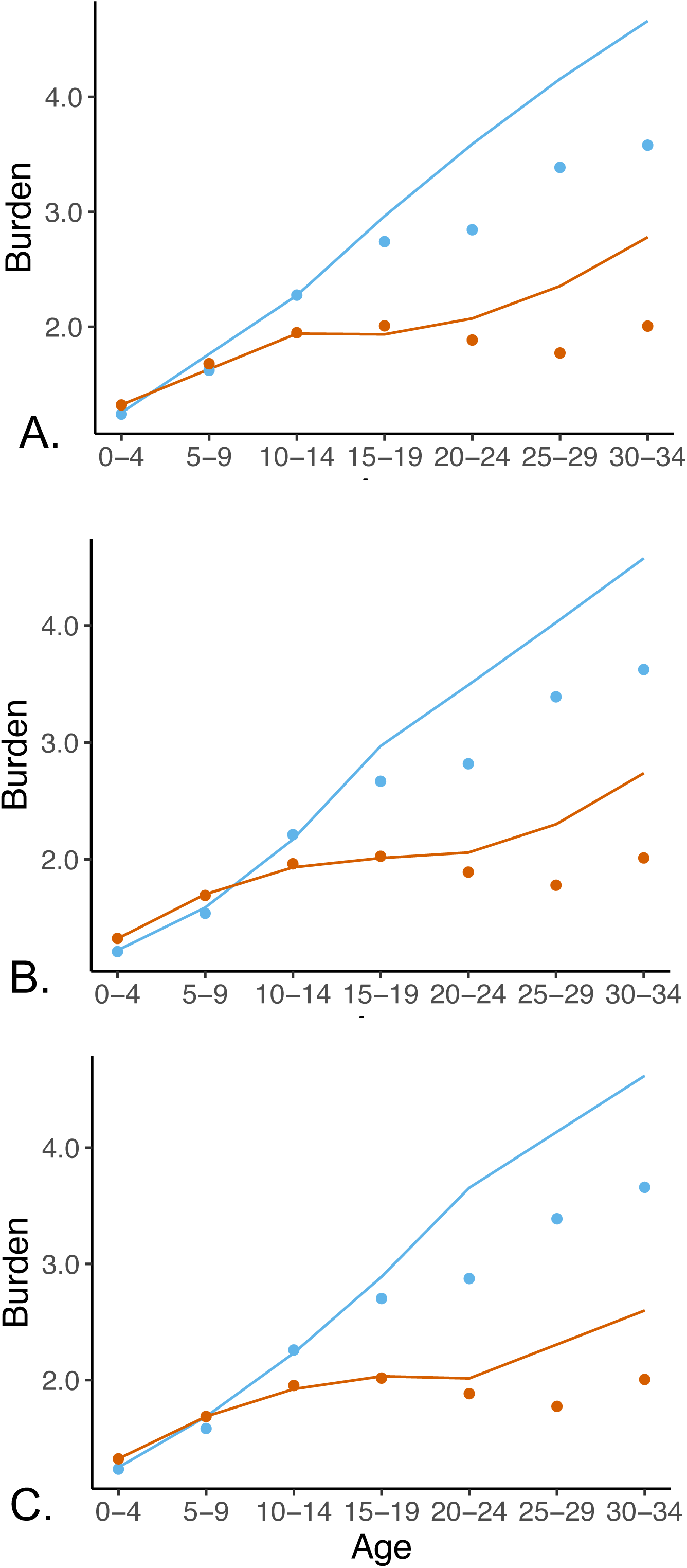

**Table 1.**
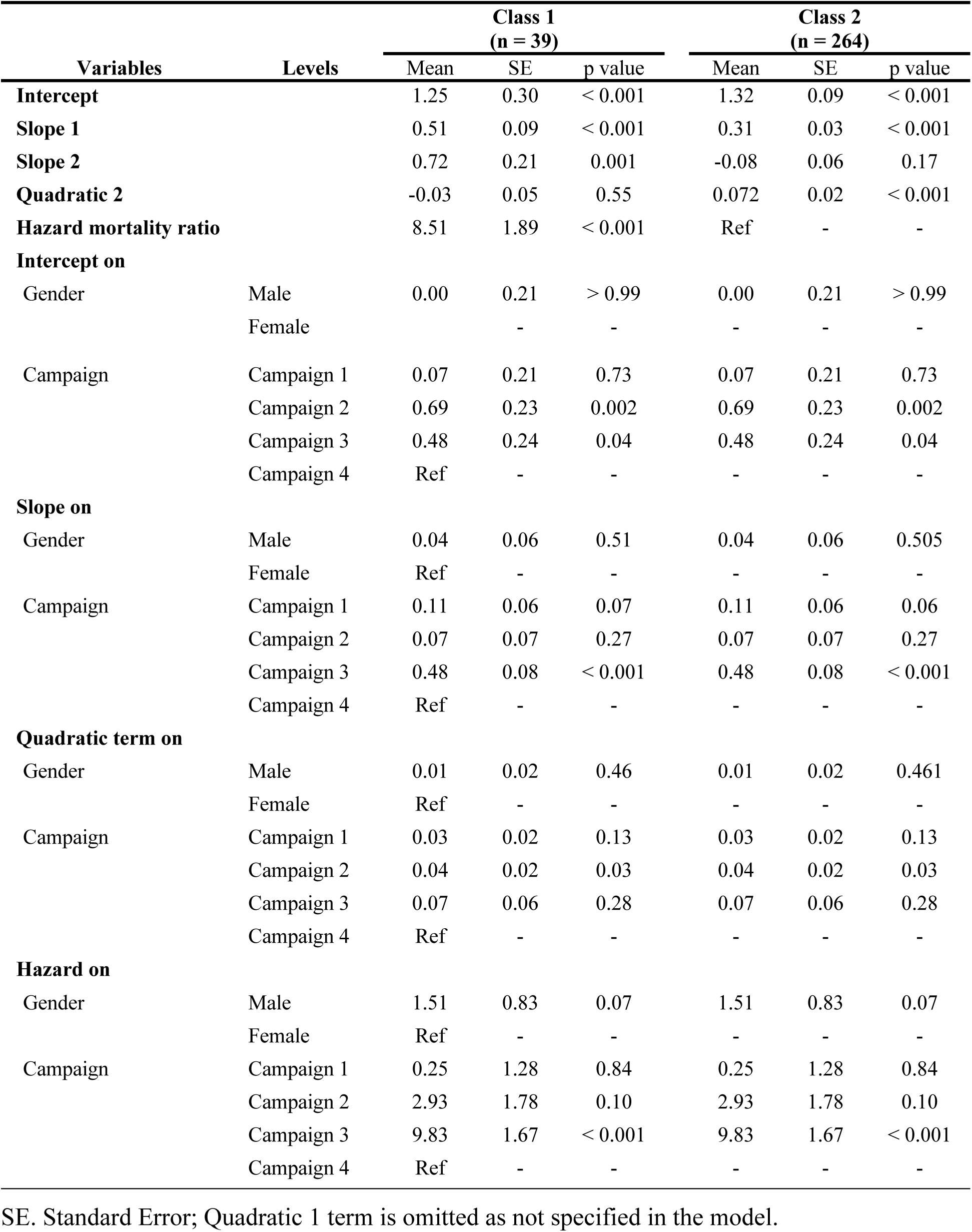
Estimates of the joint latent class model parameters.

| Variables | Levels | Class 1<br>(n = 39) |  |  | Class 2<br>(n = 264) |  |  |
| --- | --- | --- | --- | --- | --- | --- | --- |
|  |  | Mean | SE | p value | Mean | SE | p value |
| **Intercept** |  | 1.25 | 0.30 | < 0.001 | 1.32 | 0.09 | < 0.001 |
| **Slope 1** |  | 0.51 | 0.09 | < 0.001 | 0.31 | 0.03 | < 0.001 |
| **Slope 2** |  | 0.72 | 0.21 | 0.001 | -0.08 | 0.06 | 0.17 |
| **Quadratic 2** |  | -0.03 | 0.05 | 0.55 | 0.072 | 0.02 | < 0.001 |
| **Hazard mortality ratio** |  | 8.51 | 1.89 | < 0.001 | Ref | - | - |
| **Intercept on** |  |  |  |  |  |  |  |
| Gender | Male | 0.00 | 0.21 | > 0.99 | 0.00 | 0.21 | > 0.99 |
|  | Female |  | - | - | - | - | - |
| Campaign | Campaign 1 | 0.07 | 0.21 | 0.73 | 0.07 | 0.21 | 0.73 |
|  | Campaign 2 | 0.69 | 0.23 | 0.002 | 0.69 | 0.23 | 0.002 |
|  | Campaign 3 | 0.48 | 0.24 | 0.04 | 0.48 | 0.24 | 0.04 |
|  | Campaign 4 | Ref | - | - | - | - | - |
| **Slope on** |  |  |  |  |  |  |  |
| Gender | Male | 0.04 | 0.06 | 0.51 | 0.04 | 0.06 | 0.505 |
|  | Female | Ref | - | - | - | - | - |
| Campaign | Campaign 1 | 0.11 | 0.06 | 0.07 | 0.11 | 0.06 | 0.06 |
|  | Campaign 2 | 0.07 | 0.07 | 0.27 | 0.07 | 0.07 | 0.27 |
|  | Campaign 3 | 0.48 | 0.08 | < 0.001 | 0.48 | 0.08 | < 0.001 |
|  | Campaign 4 | Ref | - | - | - | - | - |
| **Quadratic term on** |  |  |  |  |  |  |  |
| Gender | Male | 0.01 | 0.02 | 0.46 | 0.01 | 0.02 | 0.461 |
|  | Female | Ref | - | - | - | - | - |
| Campaign | Campaign 1 | 0.03 | 0.02 | 0.13 | 0.03 | 0.02 | 0.13 |
|  | Campaign 2 | 0.04 | 0.02 | 0.03 | 0.04 | 0.02 | 0.03 |
|  | Campaign 3 | 0.07 | 0.06 | 0.28 | 0.07 | 0.06 | 0.28 |
|  | Campaign 4 | Ref | - | - | - | - | - |
| **Hazard on** |  |  |  |  |  |  |  |
| Gender | Male | 1.51 | 0.83 | 0.07 | 1.51 | 0.83 | 0.07 |
|  | Female | Ref | - | - | - | - | - |
| Campaign | Campaign 1 | 0.25 | 1.28 | 0.84 | 0.25 | 1.28 | 0.84 |
|  | Campaign 2 | 2.93 | 1.78 | 0.10 | 2.93 | 1.78 | 0.10 |
|  | Campaign 3 | 9.83 | 1.67 | < 0.001 | 9.83 | 1.67 | < 0.001 |
|  | Campaign 4 | Ref | - | - | - | - | - |

SE. Standard Error; Quadratic 1 term is omitted as not specified in the model.

As illustrated by the survival curves in Figure 2B, the model predicted that nearly one half of individuals in class 1 died before the age of 19, and almost no one survived later than 29 years of age. By contrast, more than 75% of individuals in class 2 were still alive at 30-34 years of age.

### 2.2. Class comparison

The sociodemographic and psychopathological characteristics of the 2 classes are presented in Table 2. As expected from the between-class discrepancy in mean age at death, we found significant differences in terms of academic level, civil status, mean number of children and household. By contrast, the sub-groups were comparable with respect to the gender ratio and the recruitment campaign. In terms of psychiatric diagnosis, the 2 classes differed only in terms of the proportion of individuals suffering from affective disorders at death, which was significantly higher in class 2 than in class 1 (57% vs 38%).

**Table 2.**
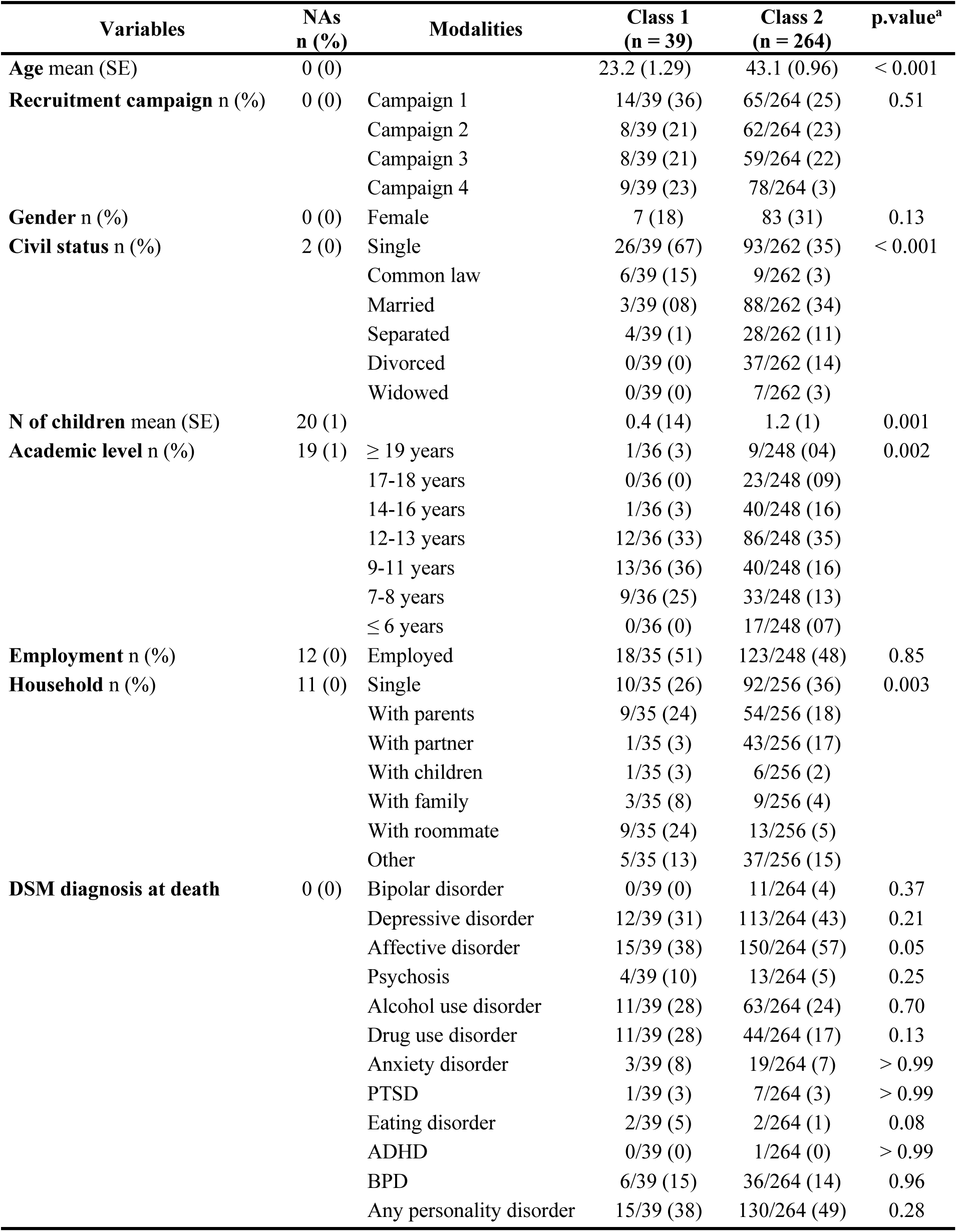
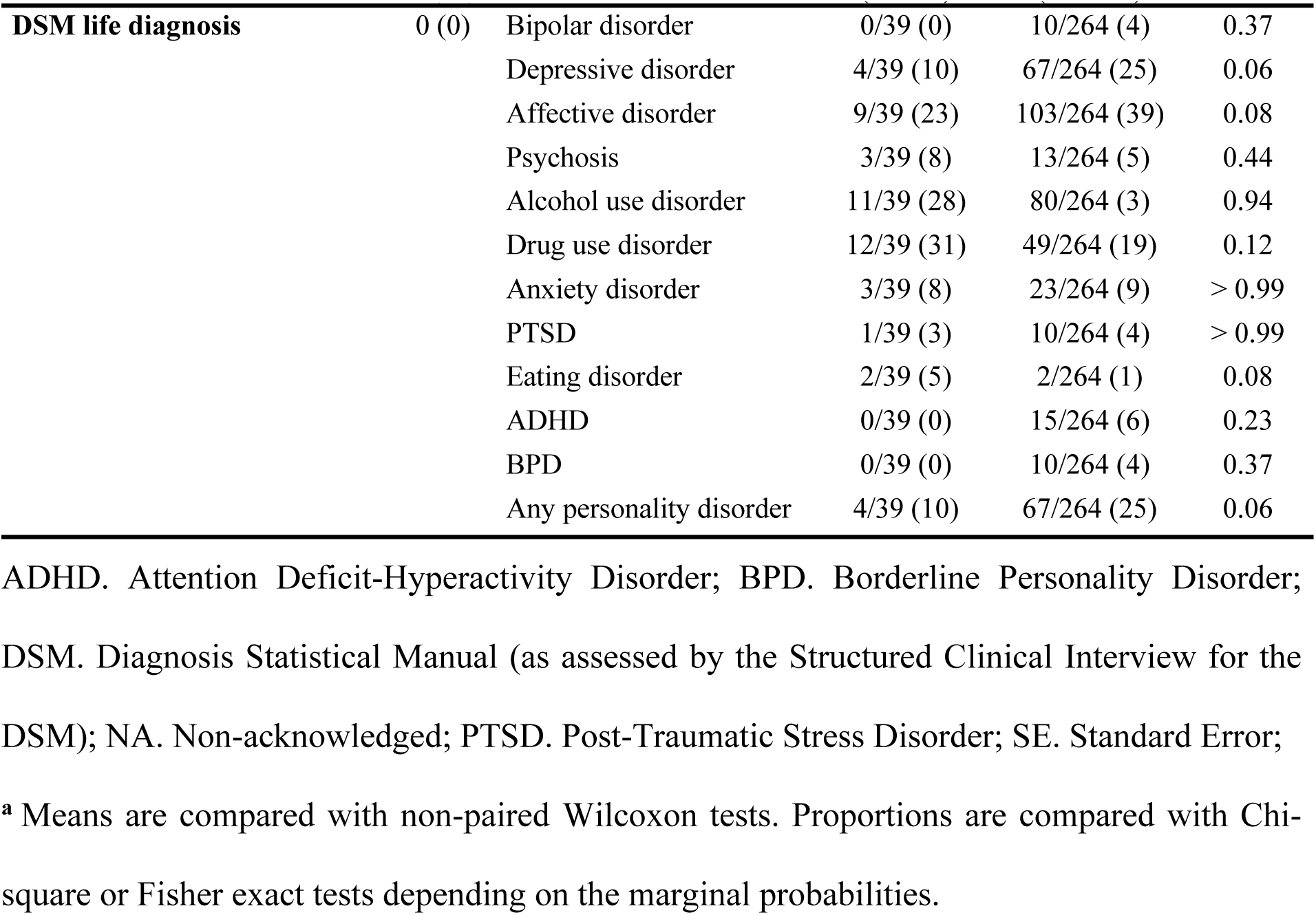
Inter-class comparison of sociodemographic and psychopathological characteristics.

| Variables | NAs<br>n (%) | Modalities | Class 1<br>(n = 39) | Class 2<br>(n = 264) | p.value<sup>a</sup> |
| --- | --- | --- | --- | --- | --- |
| **Age** mean (SE) | 0 (0) |  | 23.2 (1.29) | 43.1 (0.96) | < 0.001 |
| **Recruitment campaign** n (%) | 0 (0) | Campaign 1 | 14/39 (36) | 65/264 (25) | 0.51 |
|  |  | Campaign 2 | 8/39 (21) | 62/264 (23) |  |
|  |  | Campaign 3 | 8/39 (21) | 59/264 (22) |  |
|  |  | Campaign 4 | 9/39 (23) | 78/264 (3) |  |
| **Gender** n (%) | 0 (0) | Female | 7 (18) | 83 (31) | 0.13 |
| **Civil status** n (%) | 2 (0) | Single | 26/39 (67) | 93/262 (35) | < 0.001 |
|  |  | Common law | 6/39 (15) | 9/262 (3) |  |
|  |  | Married | 3/39 (08) | 88/262 (34) |  |
|  |  | Separated | 4/39 (1) | 28/262 (11) |  |
|  |  | Divorced | 0/39 (0) | 37/262 (14) |  |
|  |  | Widowed | 0/39 (0) | 7/262 (3) |  |
| **N of children** mean (SE) | 20 (1) |  | 0.4 (14) | 1.2 (1) | 0.001 |
| **Academic level** n (%) | 19 (1) | ≥ 19 years | 1/36 (3) | 9/248 (04) | 0.002 |
|  |  | 17-18 years | 0/36 (0) | 23/248 (09) |  |
|  |  | 14-16 years | 1/36 (3) | 40/248 (16) |  |
|  |  | 12-13 years | 12/36 (33) | 86/248 (35) |  |
|  |  | 9-11 years | 13/36 (36) | 40/248 (16) |  |
|  |  | 7-8 years | 9/36 (25) | 33/248 (13) |  |
|  |  | ≤ 6 years | 0/36 (0) | 17/248 (07) |  |
| **Employment** n (%) | 12 (0) | Employed | 18/35 (51) | 123/248 (48) | 0.85 |
| **Household** n (%) | 11 (0) | Single | 10/35 (26) | 92/256 (36) | 0.003 |
|  |  | With parents | 9/35 (24) | 54/256 (18) |  |
|  |  | With partner | 1/35 (3) | 43/256 (17) |  |
|  |  | With children | 1/35 (3) | 6/256 (2) |  |
|  |  | With family | 3/35 (8) | 9/256 (4) |  |
|  |  | With roommate | 9/35 (24) | 13/256 (5) |  |
|  |  | Other | 5/35 (13) | 37/256 (15) |  |
| **DSM diagnosis at death** | 0 (0) | Bipolar disorder | 0/39 (0) | 11/264 (4) | 0.37 |
|  |  | Depressive disorder | 12/39 (31) | 113/264 (43) | 0.21 |
|  |  | Affective disorder | 15/39 (38) | 150/264 (57) | 0.05 |
|  |  | Psychosis | 4/39 (10) | 13/264 (5) | 0.25 |
|  |  | Alcohol use disorder | 11/39 (28) | 63/264 (24) | 0.70 |
|  |  | Drug use disorder | 11/39 (28) | 44/264 (17) | 0.13 |
|  |  | Anxiety disorder | 3/39 (8) | 19/264 (7) | > 0.99 |
|  |  | PTSD | 1/39 (3) | 7/264 (3) | > 0.99 |
|  |  | Eating disorder | 2/39 (5) | 2/264 (1) | 0.08 |
|  |  | ADHD | 0/39 (0) | 1/264 (0) | > 0.99 |
|  |  | BPD | 6/39 (15) | 36/264 (14) | 0.96 |
|  |  | Any personality disorder | 15/39 (38) | 130/264 (49) | 0.28 |

Table 2 (continued). Inter-class comparison of sociodemographic and psychopathological characteristics
| Variables | NAs<br>n (%) | Modalities | Class 1<br>(n = 39) | Class 2<br>(n = 264) | p.value<sup>a</sup> |
| --- | --- | --- | --- | --- | --- |
| DSM life diagnosis | 0 (0) | Bipolar disorder | 0/39 (0) | 10/264 (4) | 0.37 |
|  |  | Depressive disorder | 4/39 (10) | 67/264 (25) | 0.06 |
|  |  | Affective disorder | 9/39 (23) | 103/264 (39) | 0.08 |
|  |  | Psychosis | 3/39 (8) | 13/264 (5) | 0.44 |
|  |  | Alcohol use disorder | 11/39 (28) | 80/264 (3) | 0.94 |
|  |  | Drug use disorder | 12/39 (31) | 49/264 (19) | 0.12 |
|  |  | Anxiety disorder | 3/39 (8) | 23/264 (9) | > 0.99 |
|  |  | PTSD | 1/39 (3) | 10/264 (4) | > 0.99 |
|  |  | Eating disorder | 2/39 (5) | 2/264 (1) | 0.08 |
|  |  | ADHD | 0/39 (0) | 15/264 (6) | 0.23 |
|  |  | BPD | 0/39 (0) | 10/264 (4) | 0.37 |
|  |  | Any personality disorder | 4/39 (10) | 67/264 (25) | 0.06 |

ADHD. Attention Deficit-Hyperactivity Disorder; BPD. Borderline Personality Disorder; DSM. Diagnosis Statistical Manual (as assessed by the Structured Clinical Interview for the DSM); NA. Non-acknowledged; PTSD. Post-Traumatic Stress Disorder; SE. Standard Error;
<sup>a</sup> Means are compared with non-paired Wilcoxon tests. Proportions are compared with Chi-square or Fisher exact tests depending on the marginal probabilities.

To probe the early factors that differentially shaped the diathesis of the two classes, we compared them according to the nature of the distal ALE occurring between ages 0 and 9 years. As observable in Table 3, left columns, the classes differed significantly in the proportion of individuals concerned by conflicts or tensions with close family members (54% in class 1 vs 31% in class 2), arrival of a new partner in one of the parents’ lives (10% in class 1 vs 2% in class 2), learning disabilities (33% in class 1 vs 18% in class 2, p = 0.018) and exposure to the suicide of a friend or family member (21% in class 1 vs 7% in class 2).

**Table 3.** Between-class comparison of the frequency of adverse events experienced in childhood (0-9), adolescence (10-14) and in the year prior to death

|  | 0-9 years old |  |  | 10-14 years old |  |  | Year prior to death |  |  |
| --- | --- | --- | --- | --- | --- | --- | --- | --- | --- |
|  | Class 1 | Class 2 | p value<sup>•</sup> | Class 1 | Class 2 | p value<sup>•</sup> | Class 1 | Class 2 | p value<sup>a</sup> |
|  | n<br>(%) | n<br>(%) |  | n<br>(%) | n<br>(%) |  | n<br>(%) | n<br>(%) |  |
| **Adversity related to the family of origin** |  |  |  |  |  |  |  |  |  |
| Victim of intra-family sexual violence | 7/39<br>(18) | 41/264<br>(16) | 0.88 | 10/39<br>(26) | 30/264<br>(11) | 0.03 | 5/39<br>(13) | 7/264<br>(0.03) | 0.01 |
| Conflicts or tensions with (a) close family member(s) | 21/39<br>(54) | 80/264<br>(0.31) | 0.004 | 24/39<br>(62) | 105/264<br>(40) | 0.01 | 19/39<br>(49) | 43/264<br>(0.16) | < 0.001 |
| Affective distance with (a) close family member(s) | 1/39<br>(3) | 17/264<br>(6) | 0.48 | 1/39<br>(3) | 19/264<br>(7) | 0.49 | 2/39<br>(5) | 9/264<br>(0.03) | 0.64 |
| Parental neglect | 13/39<br>(33) | 94/264<br>(36) | 0.92 | 15/39<br>(38) | 81/264<br>(31) | 0.43 | 9/39<br>(23) | 19/264<br>(0.07) | 0.004 |
| Inadequate education | 9/39<br>(23) | 48/264<br>(18) | 0.47 | 9/39<br>(23) | 54/264<br>(20) | 0.71 | 4/39<br>(11) | 16/264<br>(0.06) | 0.12 |
| Forced to keep or left in ignorance of a secret | 2/39<br>(5) | 20/264<br>(8) | 0.83 | 1/39<br>(3) | 27/264<br>(1) | 0.21 | 0/39<br>(00) | 12/264<br>(0.05) | 0.38 |
| Separation with (a) close family member(s) | 8/39<br>(21) | 53/264<br>(20) | > 0.99 | 4/39<br>(0) | 58/264<br>(22) | 0.14 | 3/39<br>(08) | 25/264<br>(0.09) | > 0.99 |
| Mental health issues in the family of origin | 2/39<br>(5) | 41/264<br>(16) | 0.14 | 2/39<br>(5) | 42/264<br>(16) | 0.12 | 5/39<br>(13) | 7/264<br>(0.03) | 0.01 |
| New partner in parent's life | 4/39<br>(10) | 5/264<br>(2) | 0.02 | 1/39<br>(0.03) | 2/264<br>(1) | 0.34 | 0/39<br>(0) | 1/264<br>(0.00) | > 0.99 |
| **Adversity related to affective life** |  |  |  |  |  |  |  |  |  |
| End of a romantic relationship | 0/39<br>(0) | 0/264<br>(0) | - | 3/39<br>(8) | 8/264<br>(3) | 0.16 | 8/39<br>(21) | 67/264<br>(0.25) | 0.65 |
| Tensions and/or arguments in the couple | 0/39<br>(0) | 0/264<br>(0) | - | 0/39<br>(0) | 2/264<br>(1) | > 0.99 | 0/39 (0) | 37/264<br>(0.14) | 0.007 |
| Extra-conjugal relationship of the partner | 0/39<br>(0) | 0/264<br>(0) | - | 0/39<br>(0) | 1/264<br>(0) | > 0.99 | 3/39 (8) | 11/264<br>(0.04) | 0.40 |

Table 3. Between-class comparison of the frequency of adverse events experienced in childhood (0-9), adolescence (10-14) and in the year prior to death (continued)
|  | 0-9 years old |  |  | 10-14 years old |  |  | Year prior to death |  |  |
| --- | --- | --- | --- | --- | --- | --- | --- | --- | --- |
|  | Class 1 | Class 2 | p value<sup>•</sup> | Class 1 | Class 2 | p value<sup>•</sup> | Class 1 | Class 2 | p value<sup>a</sup> |
|  | n<br>(%) | n<br>(%) |  | n<br>(%) | n<br>(%) |  | n<br>(%) | n<br>(%) |  |
| **Adversity related to children** |  |  |  |  |  |  |  |  |  |
| Mental health problem in child(ren) | 0/39<br>(0) | 0/264<br>(0) | - | 0/39<br>(0) | 0/264<br>(0) | - | 0/39 (0) | 17/264<br>(0.06) | 0.14 |
| Conflicts with child(ren) | 0/39<br>(0) | 0/264<br>(0) | - | 0/39<br>(0) | 0/264<br>(0) | - | 0/39 (0) | 25/264<br>(9) | 0.06 |
| Change in the frequency of interaction with child(ren) | 0/39<br>(0) | 0/264<br>(0) | - | 0/39<br>(0) | 0/264<br>(0) | - | 2/39 (5) | 12/264<br>(5) | 0.70 |
| **Personal adversity** |  |  |  |  |  |  |  |  |  |
| Victim of psychological violence | 7/39<br>(18) | 62/264<br>(23) | 0.57 | 8/39<br>(21) | 60/264<br>(23) | 0.92 | 5/39<br>(13) | 20/264<br>(8) | 0.34 |
| Victim of physical violence | 3/39<br>(8) | 27/264<br>(10) | 0.78 | 2/39<br>(5) | 26/264<br>(10) | 0.55 | 1/39 (3) | 1/264<br>(0) | 0.24 |
| Personal physical health problem | 5/39<br>(13) | 44/264<br>(17) | 0.65 | 9/39<br>(23) | 33/264<br>(12) | 0.08 | 10/39<br>(26) | 83/264<br>(2) | 0.46 |
| Personal psychological health problem | 6/39<br>(15) | 11/264<br>(4) | 0.44 | 9/39<br>(23) | 17/264<br>(6) | 0.44 | 6/39<br>(15) | 3/264<br>(1) | < 0.001 |
| Personal behavioral problems | 5/39<br>(13) | 18/264<br>(7) | 0.32 | 17/39<br>(44) | 45/264<br>(17) | < 0.001 | 10/39<br>(26) | 10/264<br>(4) | < 0.001 |
| Serious accident | 0/39<br>(0) | 5/264<br>(2) | > 0.99 | 0/39<br>(0) | 3/264<br>(1) | > 0.99 | 2/39<br>(5) | 18/264<br>(7) | > 0.99 |
| Witness or victim of a traumatic event | 4/39<br>(1) | 13/264<br>(5) | 0.25 | 4/39<br>(1) | 17/264<br>(6) | 0.33 | 0/39 (0) | 10/264<br>(4) | 0.37 |

Table 3. Between-class comparison of the frequency of adverse events experienced in childhood (0-9), adolescence (10-14) and in the year prior to death (continued)
|  | 0-9 years old |  |  | 10-14 years old |  |  | Year prior to death |  |  |
| --- | --- | --- | --- | --- | --- | --- | --- | --- | --- |
|  | Class 1 | Class 2 | p value<sup>•</sup> | Class 1 | Class 2 | p value<sup>•</sup> | Class 1 | Class 2 | p value<sup>a</sup> |
|  | n<br>(%) | n<br>(%) |  | n<br>(%) | n<br>(%) |  | n<br>(%) | n<br>(%) |  |
| **Adversity related to academic or professional life** |  |  |  |  |  |  |  |  |  |
| Learning disabilities, poor academic performance or school dropout | 13/39<br>(33) | 48/264<br>(18) | 0.05 | 17/39<br>(44) | 72/264<br>(27) | 0.06 | 10/39<br>(26) | 21/264<br>(8) | 0.004 |
| Relational problems with peers, bullying or harassment at school | 3/39<br>(8) | 22/264<br>(8) | > 0.99 | 7/39<br>(18) | 24/264<br>(9) | 0.09 | 2/39<br>(5) | 6/264<br>(2) | 0.28 |
| Difficulties in finding work | 0/39<br>(0) | 0/264<br>(0) | - | 0/39<br>(0) | 0/264<br>(0) | - | 3/39<br>(8) | 12/264<br>(5) | 0.42 |
| Conflicts with colleagues or boss | 0/39<br>(0) | 0/264<br>(0) | - | 0/39<br>(0) | 0/264<br>(0) | - | 0/39<br>(0) | 11/264<br>(5) | 0.37 |
| Job loss | 0/39<br>(0) | 0/264<br>(0) | - | 0/39<br>(0) | 0/264<br>(0) | - | 2/39<br>(5) | 4/264<br>(2) | 0.17 |
| Unemployment | 0/39<br>(0) | 0/264<br>(0) | - | 0/39<br>(0) | 0/264<br>(0) | - | 6/39<br>(15) | 53/264<br>(2) | 0.64 |
| **Adversity related to the extended family** |  |  |  |  |  |  |  |  |  |
| Change in the frequency of interaction with a member of the extended family | 0/39<br>(0) | 0/264<br>(0) | - | 0/39<br>(0) | 0/264<br>(0) | - | 6/39<br>(15) | 32/264<br>(12) | 0.60 |
| Relational difficulties with a member of the extended family | 0/39<br>(0) | 0/264<br>(0) | - | 0/39<br>(0) | 0/264<br>(0) | - | 2/39<br>(5) | 34/264<br>(13) | 0.20 |
| Mental health problems in a member of the extended family | 0/39<br>(0) | 0/264<br>(0) | - | 0<br>(0) | 0/264<br>(0) | - | 8/39<br>(21) | 41/264<br>(16) | 0.59 |
| Physical health problems in a member of the extended family | 0/39<br>(0) | 0/264<br>(0) | - | 0<br>(0) | 0/264<br>(0) | - | 0/39<br>(0) | 15/264<br>(6) | 0.28 |

Table 3. Between-class comparison of the frequency of adverse events experienced in childhood (0-9), adolescence (10-14) and in the year prior to death (Continued)
|  | 0-9 years old |  |  | 10-14 years old |  |  | Year prior to death |  |  |
| --- | --- | --- | --- | --- | --- | --- | --- | --- | --- |
|  | Class1<br>n<br>(%) | Class1<br>n<br>(%) | p value* | Class1<br>n<br>(%) | Class1<br>n<br>(%) | p value* | Class1<br>n<br>(%) | Class2<br>n<br>(%) | p value<sup>a</sup> |
| **Adversity related to social life** |  |  |  |  |  |  |  |  |  |
| Difficulty making friends or engaging in social relationships | 3/39<br>(8) | 20/264<br>(8) | 1.00 | 7<br>(18) | 26/264<br>(10) | 0.16 | 5/39<br>(13) | 32/264<br>(12) | 0.80 |
| Loss of (a) friend(s) | 1/39<br>(3) | 5/264<br>(2) | 0.57 | 3<br>(8) | 7/264<br>(3) | 0.13 | 2/39 (5) | 14/264<br>(5) | > 0.99 |
| Conflictual interactions with (a) friend(s) or rejection | 1/39<br>(3) | 4/264<br>(2) | 0.50 | 3<br>(8) | 5/264<br>(2) | 0.07 | 9/39<br>(24) | 28/264<br>(11) | 0.04 |
| Social isolation | 3/39<br>(8) | 15/264<br>(6) | 0.71 | 9<br>(23) | 17/264<br>(6) | 0.002 | 7/39<br>(18) | 73/264<br>(28) | 0.28 |
| **Adversity related to losses** |  |  |  |  |  |  |  |  |  |
| Move or departure | 10/39<br>(26) | 42/264<br>(16) | 0.13 | 7<br>(18) | 22/264<br>(8) | 0.08 | 9/39<br>(24) | 34/264<br>(13) | 0.14 |
| Death of a close relative | 1/39<br>(3) | 6/264<br>(2) | > 0.99 | 1<br>(3) | 12/264<br>(5) | > 0.99 | 2/39<br>(5) | 8/264<br>(3) | 0.62 |
| Suicide attempt of a relative | 1/39<br>(3) | 5/264<br>(2) | 0.57 | 3<br>(8) | 14/264<br>(5) | 0.47 | 1/39 (3) | 12/264<br>(5) | > 0.99 |
| Suicide of a relative | 8/39<br>(21) | 19/264<br>(7) | 0.01 | 11<br>(28) | 27/264<br>(10) | 0.004 | 3/39<br>(8) | 22/264<br>(8) | > 0.99 |
| **Adversity related to living conditions** |  |  |  |  |  |  |  |  |  |
| Precarious living conditions | 0/39<br>(0) | 17/264<br>(6) | 0.14 | 0<br>(0) | 14/264<br>(5) | 0.23 | 2/39 (5) | 20/264<br>(7) | > 0.99 |
| Homelessness | 0/39<br>(0) | 0/264<br>(0) | - | 0<br>(0) | 1/264<br>(0) | > 0.99 | 2/39<br>(5) | 12/264<br>(5) | 0.70 |
| Financial problems | 0/39<br>(0) | 3/264<br>(1) | > 0.99 | 0<br>(0) | 4/264<br>(2) | > 0.99 | 9/39<br>(23) | 91/264<br>(34) | 0.22 |
| Legal proceedings against the individual | 0/39<br>(0) | 0/264<br>(0) | - | 1<br>(3) | 4/264<br>(2) | 0.50 | 4/39<br>(10) | 18/264<br>(7) | 0.50 |

Only relevant events with a frequency equal to or greater than 5% in at least one class and one of the periods under consideration are reported.
<sup>a</sup> Comparisons are performed with Chi-square or Fisher exact tests depending on the marginal probabilities.

We then focused on the 10-14 period to examine the ALE that occurred simultaneously with the bifurcation of the trajectories (Table 3, middle columns). The proportion of individuals who experienced social isolation or conflicts within the family unit during early adolescence was significantly higher in class 1 than in class 2 (44% vs 17%, and 62% vs 40%, respectively). The two classes also differed significantly in the proportion of members exposed to intra-family sexual violence (26% in class 1 vs 11% in class 2) or suicide of a relative (28% in class 1 vs 10% in class 2) between 10 and 14. Finally, we found significantly more frequent behavioral problems in class 1 than in class 2 at these ages (44% vs 17%).

With respect to proximal ALE (see Table 3, right columns), more class 1 individuals had experienced the following events within the year prior to death, compared with class 2 individuals: conflicts or tensions with a close family member (49% in class 1 vs 16% in class 2), parental neglect (23% in class 1 vs 7% in class 2) or conflict with a friend (24% in class 1 vs 11% in class 2). In addition, there were more frequent behavioral and/or psychological problems in class 1 than in class 2 at the time of death (26% vs 4% and 15% vs 1%, respectively). Conversely, a higher proportion of class 2 individuals was exposed to conjugal tensions or arguments in their last year of life (14% vs 0%).

## 3. Discussion

In this paper, we relied on a mixed quantitative/qualitative approach to specify how adolescence influences the evolution of the stress-diathesis interplay in individuals who died by suicide. We found that 2 subpopulations were relevantly distinguishable as regards a common underlying developmental process.

While both classes had experienced growing adversity during infancy, the divergence of the trajectories increased as of early adolescence. For individuals dying at middle age (class 2), the allostatic regulation appeared to stabilize temporarily during young adulthood before progressively faltering again until death. By contrast, suicides in late adolescence (class 1) were preceded by a constant increase in BA, suggesting a progressive acceleration in the attrition of the allostatic mechanisms. Interestingly, both distal and proximal portions of the adolescent trajectories were characterized by the overrepresentation of dependent adverse events, i.e. events that likely occurred non-randomly due to the individual/environment interrelationship^20^ (e.g. school difficulties, academic dropouts, conflicts with relatives). There have been several observations in the developmental literature that maladaptive response to stress may cause more frequent stressful events to occur^27–29^. The process might be particularly prominent during adolescence, as studies have provided clues that teenagers subjected to stressful conditions may be more prone to risk-taking and novelty-seeking behaviors^30^. In early-suicide trajectories, the repetition of stress-induced – stress-inducing events may have caused a vicious spiral of homeostatic perturbation revealing in age-specific contexts and reaching its critical point at young adult ages. Importantly, most of these events had a social valence, which is meaningful as adolescence is a period of shift in interests and security basis from family to peers. Both distally and proximally, the conjunction of family conflicts and lack of extra-family support may have played a prominent role in adolescents’ allostatic failure by thwarting a sense of social belongingness under construction^31^.

We see two compatible hypotheses to account for the bifurcation at 10-14. First, a substantial proportion of the 2 subgroups experienced early ALE that have been identified as strong predictors of suicidal behaviors, including childhood neglect, psychological, physical and sexual violence^32–35^ and health problems^36^. However, individuals who died as young adults differed significantly in terms of frequency of tensions in the family and arrival of a new partner in one of the parents’ lives. Household instability and family stress have been demonstrated to compromise internal safety, emotional regulation or conflict resolution^37^, thus altering children’s coping abilities^38^. The divergence of the 2 trajectories during adolescence could be interpreted as the delayed revelation of an early-acquired adaptive deficit due to a precarious family context. This hypothesis is consistent with repeated evidence that some neuro-biological consequences of early exposure to stress become evident during adolescence^39^. From this perspective, the break predicted by our model could be the signature of a stress-sensitivity potentiating/incubation effect that adolescence reveals^39^. Alternatively, the steady increase in BA observed in class 1 could have resulted from some ALE occurring more frequently during the 10-14 period. Biological research has provided strong evidence that adolescence is a period of critical vulnerability to adversity^40–42^, due to immature regulation of hormonal responses^43,44^ and greater stress-sensitivity of key brain regions^39,42^. There are also clues that stressful events occurring during adolescence durably compromise brain function and structure^41,42^. From this perspective, the bifurcation of the trajectories predicted by our model could reflect the plasticity of young adolescents’ allostatic system, of which extreme vulnerability to stress and possibly long-term consequences are corollaries.

Almost 25% of individuals who died as young adults (vs 10% in class 2) were victims of intrafamily sexual violence between ages 10 and 14. The literature presents robust evidence that childhood sexual assault pervasively undermines the mental health equilibrium^45^ and leads to higher risk of suicidal outcomes^32,33^. Much scarcer is the evidence about the consequences of intrafamily sexual violence occurring *during* adolescence. Yet, our results suggest that the deleterious effects of sexual victimization during puberty may differ from those of childhood abuse. In addition, 20% and 30% of individuals in class 1 (vs 7% and 10% in class 2) were exposed to the suicide of a close relative during infancy and young adolescence, respectively. Beyond the genetic and epigenetic substrates and/or shared adverse living conditions that may be implied, such aggregation raises questions about a possible suicidal contagion process. In the past 3 decades, researchers have provided epidemiological^46^ and experimental^47^ evidence that exposure to a suicide model may contribute to precipitating suicidal behaviors in vulnerable individuals. In addition to this “trigger” effect, some population-based studies have evidenced that suicide contagion mechanisms may appear up to several years after the index death^48^ through a “sleeper effect”. In our study, the role of contagion as a trigger for suicide is worth considering, as almost 10% of our whole sample was exposed to the suicide of a close relative in the year prior to their death. However, the fact that the 2 classes differed in terms of suicide exposure during infancy and adolescence indicates that suicide contagion may also result from a long term “kindling” effect, possibly through implicit encryption and retention of the suicide model^49^.

To our knowledge, this study is one of the largest psychological autopsy investigations with in-depth collection of ALE that have been carried out so far. It sheds new light on the transitional process characterizing adolescence. However, several limitations need to be taken into account. First, a common concern about retrospective collection of ALE relates to information or recall biases. In our study, we adopted proven measures to minimize such biases: systematic semi-structured explorations, conversational-style interviews, use of memory anchors, memory-stimulation techniques and cross-checking of collected data from various sources^50^. Although these precautions may not have fully compensated for the inaccuracies of subjective reporting procedures, it should be noted that we focused mainly on major ALE, which are less susceptible to oblivion, and that information biases were probably comparable for the 2 classes. A second limitation is the absence of a comparison group, which prevents us from drawing any conclusion about the role of adolescence in determining whether a person follows a suicidal trajectory or not. This highly relevant question was beyond our scope. It would deserve specific developmental investigations, possibly with the same approach as we used here. Finally, the expert rating procedure implied that BA values were available for 5-year periods. This time scale precluded precise examination of the developmental changes that occurred within the adolescence time slot. In future studies, researchers could consider developing alternative methods for transforming ALE data in order to gain in temporal acuity.

## Conclusion

Our results support the concept of adolescence as a critical developmental period through validation in the suicidal process framework. Related to the intense allostatic maturation processes that come with puberty, we predicted that young adolescence would be associated with deviations of adversity trajectories towards suicide while older adolescents’ suicide would be preceded by a specific allostatic exhaustion pattern. These observations could inform clinicians’ practices as they provide evidence for enriching assessments of patients’ suicidal risk and related services with the appraisal of the adversity sequences that they experienced, especially during adolescence. In addition, they shed light on the relevance of both holistic and dynamic approaches as the principles of an emerging and promising research paradigm for suicide. Provided that adapted methods are developed and tested, such paradigm could open whole new paths of exploration of suicidal behaviors in young people that traditional epidemiology have overlooked so far.

## Acknowledgment

We thank the Fonds de recherche du Québec Société Culture and the Fonds de recherche du Québec-Santé that financially supported the data gathering over the years, the Réseau québécois sur le suicide, les troubles de l’humeur et les troubles associés that helped for conducting the study and the CHU de Lille that granted financial and human resources to analyze the data.

