## Supplemental Material for "Life trajectories towards suicide: developmental role and specificities of adolescence"

**Supplement. Model selection**

We selected the most appropriate JLCM model according to the following decision-making process. First, we discarded models with 3 and 4 classes, due to underspecification or overfitting with regard to our data. We then compared the factorial variations of the remaining 2-class JLCM in terms of fit indices. As illustrated in eFigure 1, related patterns were similar for both BIC and AIC values. Models including quadratic curves systematically outperformed their linear counterparts, with BIC differences ranging from 40 to 214 and AIC differences ranging from 77 to 292 (for smooth and broken trajectories at 10-15 years, respectively). It is also clear from eFigure 1 that the models with broken curves fitted the data much better than the smooth-curve models. However, when the focus was on the piecewise LCGM, it was not obvious from the fit indices which of the models with a break at the 10- to 14-, 15- to 19- or 20- to 24-year period performed best. We thus plotted the respective predictions of these piecewise model variants to verify the clinical relevance of the corresponding trajectories (see eFigure 2). We evidenced that only the model with a break at 10-14 fully captured a split-point between the 2 trajectories occurring during early adolescence. We thus retained the 2-class, quadratic piecewise LCGM with a break at 10-14 years of age as best accounting for our data. For this model, AIC was 4822.4, BIC was 4993.3 and entropy was 0.91.

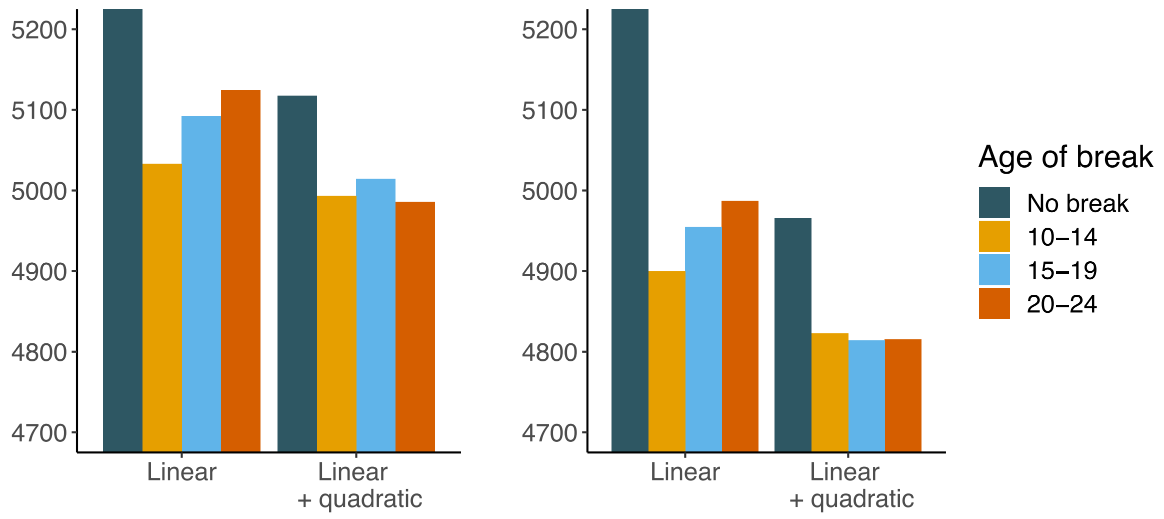

**Figure S1. Fit indices of the joint latent class models according to the curve parameters and the place of the breakpoint.** A. Bayesian Information Criterion (BIC). B. Akaike Information Criterion (AIC)

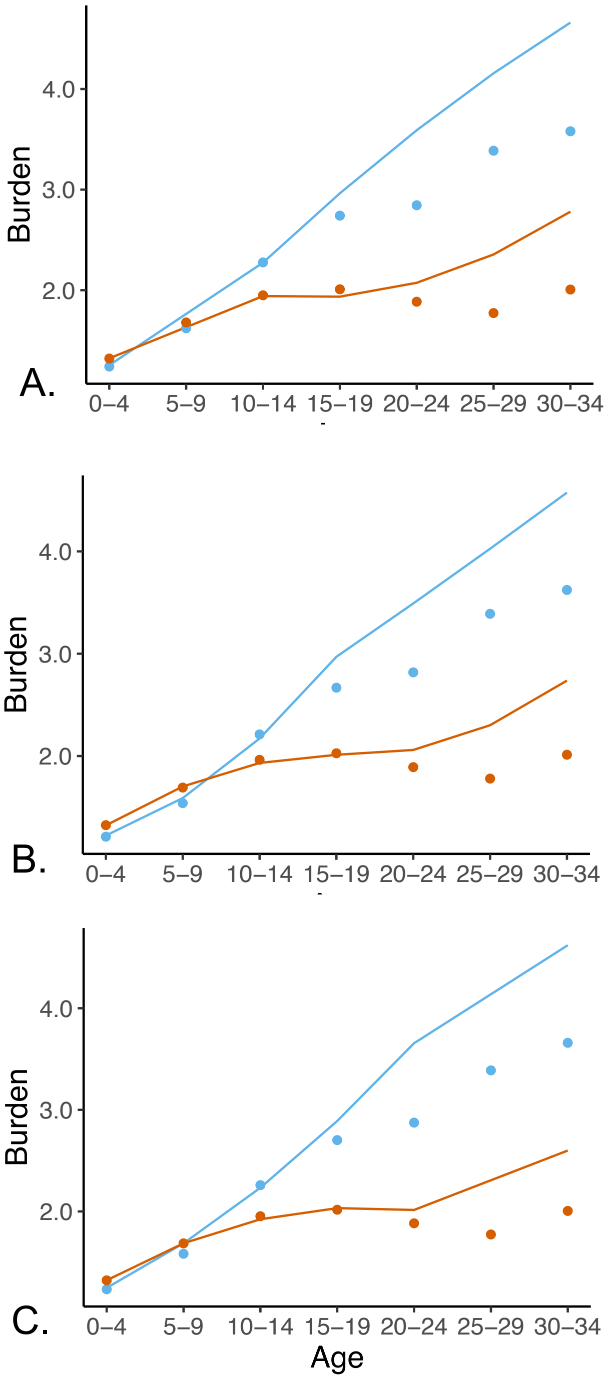

**Figure S2. Predicted and observed burden of adversity trajectories according to the JLCM variants.** The panels correspond to models where the breakpoint is fixed at A. age 10-14 years, B. age 15-19 years and C. age 20-24 years. Solid lines and dots represent the predicted and observed mean values, respectively. Class 1 trajectories are in blue, class 2 trajectories are in red.
